# A kidney specific mouse model to study the effects of *in vivo* induction of Yamanaka factors

**DOI:** 10.1101/2025.08.30.672899

**Authors:** Katharina Lemberg, Gijs A.C. Franken, Korbinian M. Riedhammer, Selina Hölzel, Kirollos Yousef, Kraisoon Lomjansook, Gina Kalkar, Caroline M. Kolvenbach, Daniel Marchuk, Elena Zion, Ken Saida, Florian Buerger, Friedhelm Hildebrandt

## Abstract

**Introduction:** Maladaptive repair after acute kidney injury (AKI) leads to fibrosis and chronic kidney disease (CKD). Improving the resilience and stimulating tissue repair after injury is crucial to prevent AKI-to-CKD transition. Using a combination of transcription factors (Yamanaka factors *Oct4, Klf4* and *Sox2, “OKS”*) to partially reprogram tissues and enhance regeneration *in vivo*, could be a promising approach as shown by amelioration after various organ injury, yet not investigated for AKI to date.

**Methods:** We used a ubiquitously and kidney-specific transgenic mouse model to investigate *OKS* expression in kidney. In a kidney-specific model using Pax8-Cre, we then induced AKI via aristolochic acid (AA), simultaneously expressing *OKS* to determine potential protective effects after kidney injury.

**Results:** We show that a ubiquitously expressing *OKS*-mouse model was not suitable due to toxic effects and limited kidney expression. In the *Pax8*-Cre mouse model, we observed expression almost exclusively to proximal tubules. While induction for more than 3 days caused dysplastic tumor formation, an induction regimen limited to 3 days was not able to improve phenotypic outcome after AA-injury.

**Conclusion:** Partial reprogramming of the kidney using OKS is feasible; however, it requires a delicate balance to the risk of oncogenic transformation. Determine a dose that effectively promotes repair without crossing the threshold into harmful effects remains a major challenge, posing significant safety concerns for translating the approach to humans in the near future.

## INTRODUCTION

Acute kidney injury (AKI) is defined by a rapid reduction of kidney function, evaluated by an increase of serum creatinine and a concurrent decrease in urine production.^1^ The etiology of AKI is diverse, comprising kidney-specific (e.g. acute glomerular, vascular or tubulointerstitial diseases), as well as extrarenal (e.g. postrenal obstruction) and systemic causes (e.g. hypovolemia, infections, nephrotoxic drugs). AKI represents one of the most common conditions in hospitalized patients, affecting approximately 20% of patients,^2,3^ and increasing in more than 50% of patients in intensive care units.^4^ Moreover, AKI is concomitant with an increased risk of developing chronic kidney disease (CKD), end-stage kidney disease (ESKD), and mortality. Treatment of AKI may include discontinuation of all nephrotoxic agents, balancing of volume status and hemodynamic monitoring, and potentially kidney replacement therapy.^1^ Still, patients have an approximate 9-fold increased risk to develop CKD, underlining the need for innovative and improved treatment options. While the capacity to regenerate is very limited in glomeruli, tubular cells have the ability to proliferate after dedifferentiation, replacing the irreversibly injured cells and thereby restoring tubular function.

^5^ However, AKI can also lead to fibrosis, if—after the acute phase of AKI, in which cell death occurs—the recovery phase is not successfully resolved. Various mechanisms have been identified that contribute to incomplete or maladaptive repair, with persistent tubulointerstitial inflammation, fibroblast expansion and extracellular matrix deposits.^6^ These mechanisms include partial epithelial–mesenchymal transition (EMT), emerging cell senescence and G2/M cell-cycle-arrest (CCA), as well as fibroblast and immune cell activation, and the incidence of a profibrogenic secretome.^7^ Improving the resilience and repair capacity of the proximal tubules and preventing dysfunctional repair mechanisms is therefore considered a viable target to prevent or decrease the development of AKI.

One innovative treatment approach has developed over the last decade and is based on transient expression of a cocktail of transcription factors, which, acting in concert, have shown the ability to induce pluripotency in somatic cells. These seminal studies revolutionized stem cell research^8,9^ and the original four transcription factors *OCT4, KLF4, SOX2* and *c-MYC* have been termed the Yamanaka transcription factors (YF). Since then, extensive efforts have focused on applying this pivotal concept to living animals, driven by two main expectations. Firstly, *in vivo* reprogramming tools are expected to elucidate the complex microenvironment regulating cellular (de-)differentiation within healthy and diseased tissues. Secondly, if applied with strict spatial and temporal control, strategies to (de-)differentiate somatic cells directly within the organism could open up novel and clinically relevant approaches in regenerative medicine.^10^ Multiple studies have so far investigated effects of this *in vivo* approach, mostly using transgenic mouse models,^11–19^ but also viral vectors,^20–22^ or plasmids to target tissues. ^23^ Applications of YF expression in a disease context with beneficial impact have been very broad: the majority of studies investigated aging-related changes, showing rejuvenation of different organs, extended life span and behavioral improvements.^11,12,17–19,22,24^ Many studies focused on central nervous systems and neurons, ranging from optic nerve lesions,^21^ Alzheimer disease,^25^ to traumatic or ischemic brain injury.^20,26^ Nevertheless, an alleviation after injury with less fibrosis and better outcome could be shown also for investigations involving skin,^13^ cardiomyocytes,^14^ skeletal muscle,^15,23^ and liver.^16^ This suggests that temporarily expressing YF could stimulate tissue repair or regeneration via partial dedifferentiation of the somatic cells, reduction of senescence, and the accompanying proinflammatory secretome, as well as creating a tissue-protective environment. Given this, the potential rejuvenative influence/effect of YF in the kidney could be potentially used to alleviate AKI-phenotypes.

In this study, we describe two mouse models that express 3 of the 4 initial Yamanaka factors (*Oct4, Klf4, Sox2*) within the kidney and investigate their spatial expression pattern and assess their potential for phenotypic improvement upon kidney injury.

## METHODS

### Mouse breeding, maintenance and general procedures

All experimental protocols were reviewed and approved by the Institutional Animal Care and Use Committee at Boston Children’s Hospital (#00002435). Mice were housed and handled according to the Guidelines for the Care and Use of Laboratory Animals with unlimited access to water and rodent chow, a light period of 12 hours per day and pathogen-free conditions. Animal experiments were conducted in accordance with the ARRIVE guidelines. Euthanization was performed by CO_2_ administration.

### Mouse models

We obtained tetOP-OKSmC mice (Jax Strain #:034917) which harbor two transgenes (tetO-*OKSmCherry*^tg/-^, ROSA26-*rtTA*^tg/-^): 1.) they express the three Yamanaka transcription factors *Oct4, Klf4* and *Sox2*, followed by the *mCherry* sequence, in a polycistronic construct, under the control of a tet-responsive element (*tetO*) with CMV minimal enhancer-less promoter, 2) at the ROSA26-locus these mice constitutively express the reverse tetracycline-controlled transactivator protein (rtTA).

For the kidney tubule specific expression of *OKS*, OKMSCh250 mice (Jax Strain #:031012), as well as Pax8^cre^ (Strain #:028196 ^27^) were obtained, and crossed to obtain transgenic mice of the following genotypes: tetO-*OKSmCherry*^tg/-^; *rtTA*^fl/-^; *Pax8Cre*^tg/wt^ (hereafter termed Pax8Cre^+^ mice) and tetO-*OKSmCherry*^tg/-^; *rtTA*^fl/-^; *Pax8Cre*^wt/wt^ (Pax8Cre^-^).

We used following primers to perform genotyping:

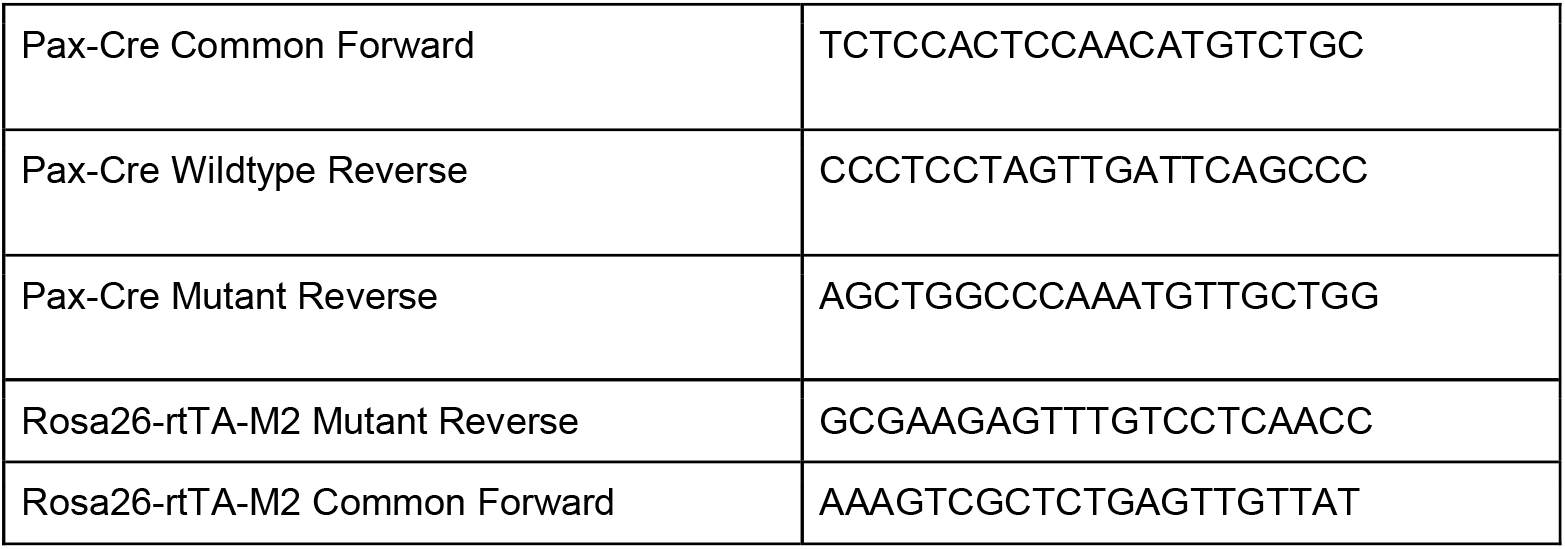
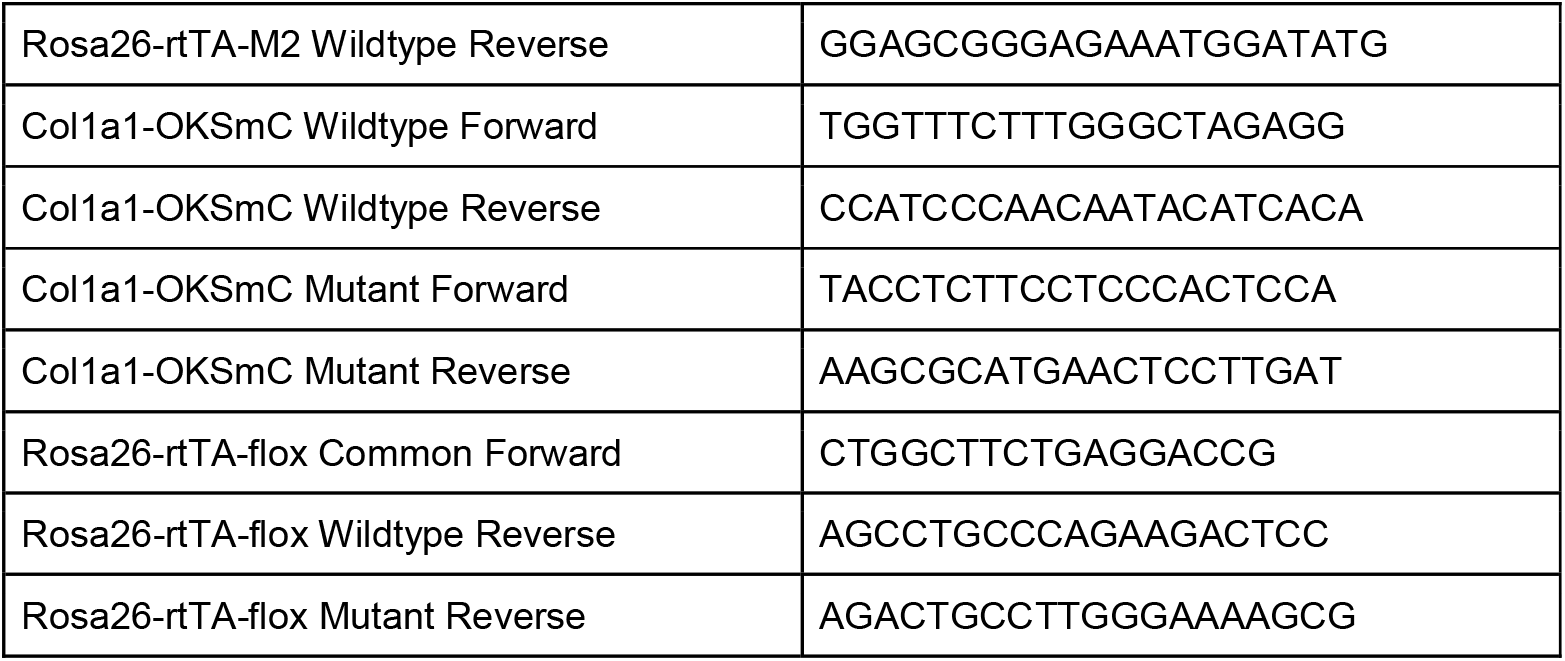

PCR Program for all genotypes was as follows:

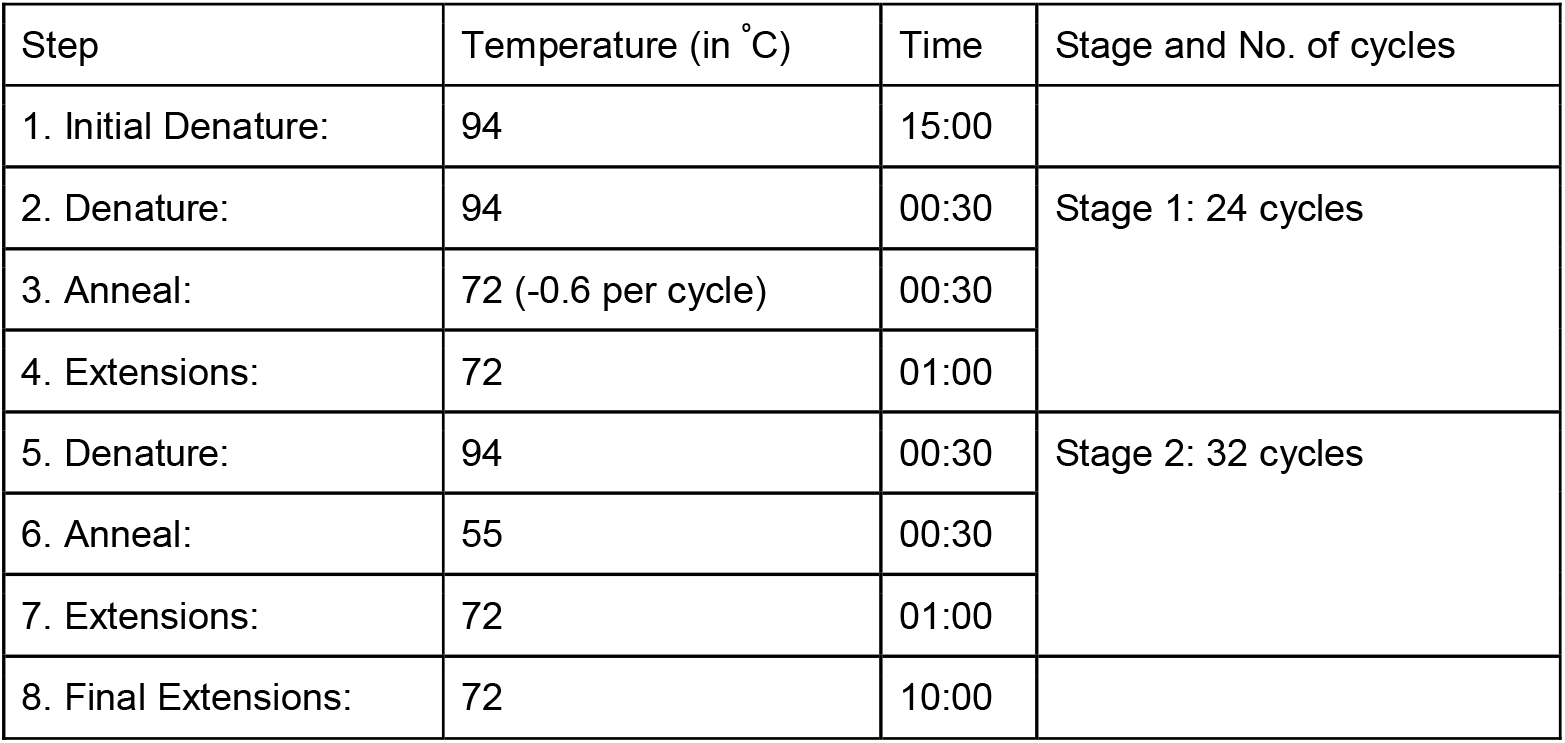

| Step | Temperature (in °C) | Time | Stage and No. of cycles |
| --- | --- | --- | --- |
| 1. Initial Denature: | 94 | 15:00 |  |
| 2. Denature: | 94 | 00:30 | Stage 1: 24 cycles |
| 3. Anneal: | 72 (-0.6 per cycle) | 00:30 |  |
| 4. Extensions: | 72 | 01:00 |  |
| 5. Denature: | 94 | 00:30 | Stage 2: 32 cycles |
| 6. Anneal: | 55 | 00:30 |  |
| 7. Extensions: | 72 | 01:00 |  |
| 8. Final Extensions: | 72 | 10:00 |  |

### OKS induction via doxycycline

OKS induction was achieved by administering doxycycline (dox; MP Biomedicals, Irvine, CA, USA) in drinking water (concentration of 1 mg/ml or 2 mg/ml, as indicated), supplemented by 5% (w/v) sucrose. Induction length ranged from 1 day to a maximum of 9 days in the induction testing experiments. Mice were monitored for weight loss and sacrificed immediately at specific time points as indicated.

### Combined kidney injury via aristolochic acid and OKS induction

In a randomized mouse cohort, we administered aristolochic acid I (AA, Sigma Aldrich, St. Louis, MO, USA) intraperitoneally in 8-21 week old male mice, diluting AA with DMSO to reach a final volume of 200 μl and a concentration of 7.5 mg/kg body weight. In the combined induction and injury experiments, mice received dox 1 mg/ml in their drinking water for 3 days in total, with administration of AA 48 hours after start of dox induction. Mice were monitored daily for weight loss or other abnormalities and euthanization was conducted after 28 days.

### Blood analysis

During dissection, 100-150 μl of blood was drawn by cardiac puncture. Blood samples were collected in lithium heparin tubes and directly analyzed by the Vetscan VS2 Chemistry Analyzer using Comprehensive Diagnostic Profile rotors.

### Immunofluorescence and Histological Studies

Mouse kidneys were sectioned in the coronal plane and tissues were incubated in paraformaldehyde, diluted in 1x PBS at a final concentration of 4% (w/v), (Electron Microscopy Sciences, Hatfield, PA, USA) overnight at 4°C on a roller bank followed by paraffinization and sectioning by the Histology core at Boston Children’s Hospital, Boston, USA. Five-micrometer-thick sections were subsequently deparaffinized and rehydrated followed by heat-induced epitope retrieval in 1x Tris-EDTA solution at pH 9 (Abcam, Waltham, MA, USA). Sections were subsequently blocked in 1% (v/v) bovine serum albumin (ZellBio GmbH, Lonsee, Germany) blocking buffer at RT for 30 min. Thereafter, sections were probed with primary antibody diluted in blocking buffer and incubated overnight at 4°C, following washing in 1x PBS the next day. Sections were then incubated in blocking buffer supplemented with corresponding secondary antibodies for 1 hour at RT. Following counterstaining with 1:5000 (v/v) DAPI (ThermoFisher, Waltham, MA, USA), samples were mounted using Fluoromount-G (ThermoFisher). Images were acquired on the Nikon-Ti2 Eclipse. Fiji version 2.17.0 was used to prepare images for visualization, Qupath-05.01 was used for the quantification using the unaltered, original acquisition.nd2 files.

**Table 2:** Antibodies used and their dilutions for immunofluorescence.

| Antibody (raised species) | Dilution (v/v) | Cat. No, company |
| --- | --- | --- |
| Sox2 (Rabbit) | 1:200 | PA1-094, ThermoFisher |
| mCherry/tdTomato (Goat) | 1:200 | AB8181-200, Origene |
| Megalin (Rat) | 1:200 | 31012, BiCell Scientific |
| Alexa Fluor 488 anti-rat IgG (Donkey) | 1:300 | A21208, ThermoFisher |
| Alexa Fluor 647 anti-rat IgG (Donkey) | 1:300 | A78947, ThermoFisher |
| Alexa Fluor 594 anti-goat IgG (Donkey) | 1:300 | A11058, ThermoFisher |
| Alexa Fluor 488 anti-rabbit IgG (Donkey) | 1:300 | A21206, ThermoFisher |
| Alexa Fluor 647 anti-rabbit IgG (Donkey) | 1:300 | A31573, ThermoFisher |

For histological studies, sections of 5 µm were stained with Masson’s trichrome (MT) and periodic acid-Schiff (PAS), following standard protocols. Whole section imaging was performed using a high-resolution digital slide scanner (SLIDEVIEW VS200, Olympus, Tokyo, Japan) at 10x magnification and acquired images were processed with the analysis software Olympus OlyVIA 3.2.1.

### Quantitative PCR

Kidney and Liver tissue samples were snap frozen in liquid nitrogen upon dissection. RNA extraction was conducted using the RNEasy Mini Kit (Qiagen). In brief, 20-30 mg of tissue was added to 500 µl of RLT Buffer, homogenized for 10 s using Tissue Tearor II (BioSpec Products, Bartlesville, OK, USA), then 70% ethanol was added and, after through mixing, the solution transferred to the provided columns. Extraction was performed following the kit protocol. After quantification by Nanodrop spectrometer (ThermoFisher) cDNA was synthesized using the iScript™ cDNA Synthesis Kit (BioRad, Hercules, CA, USA). qPCR was performed using the TaqMan™ system, using a StepOnePlus Real time machine (Thermofisher). Used probes are listed below.

### TaqMan™ probes

*Sox2*: Mm00488369_s1

*Oct4*: Mm00658129_gH

*Klf4: Mm00516104_m1*

*Mplum*: Mr07319439_mr

*B-actin*: Mm00607939_s1

*P21*: Mm04205640_g1

*Hacvr*: Mm00506686_m1

*Lcn2*: Mm01324470_m1

*P16*: Mm00494449_m1

*Col1a1*: Mm00801666_g1

### Statistics

For statistical testing Graphpad Prism 10.1.1 and R-4.5.1 were used. Statistical significance was determined using a linear mixed-effect model for weight changes over time, an unpaired Student’s T-test or a two-way ANOVA with Tukey post-hoc for gene expression tests and a standard confidence interval of 95%.

## RESULTS

### Ubiquitous tetOP-OKSmC mice transgenic mice show low renal expression for *OKS*

To investigate the effects of *in vivo* expression of *OKS*, we used mice that carried *Oct4, Klf4, Sox2*, and *mCherry* polycistronic transgenes under the control of a tetracycline responsive element (TRE), as well as a transgene at the *ROSA26* locus expressing rtTA (**Fig. 1A**). Dox exposure would activate the rtTA, thereby enabling expression of *OKS* and *mCherry*.

**Figure 1:**
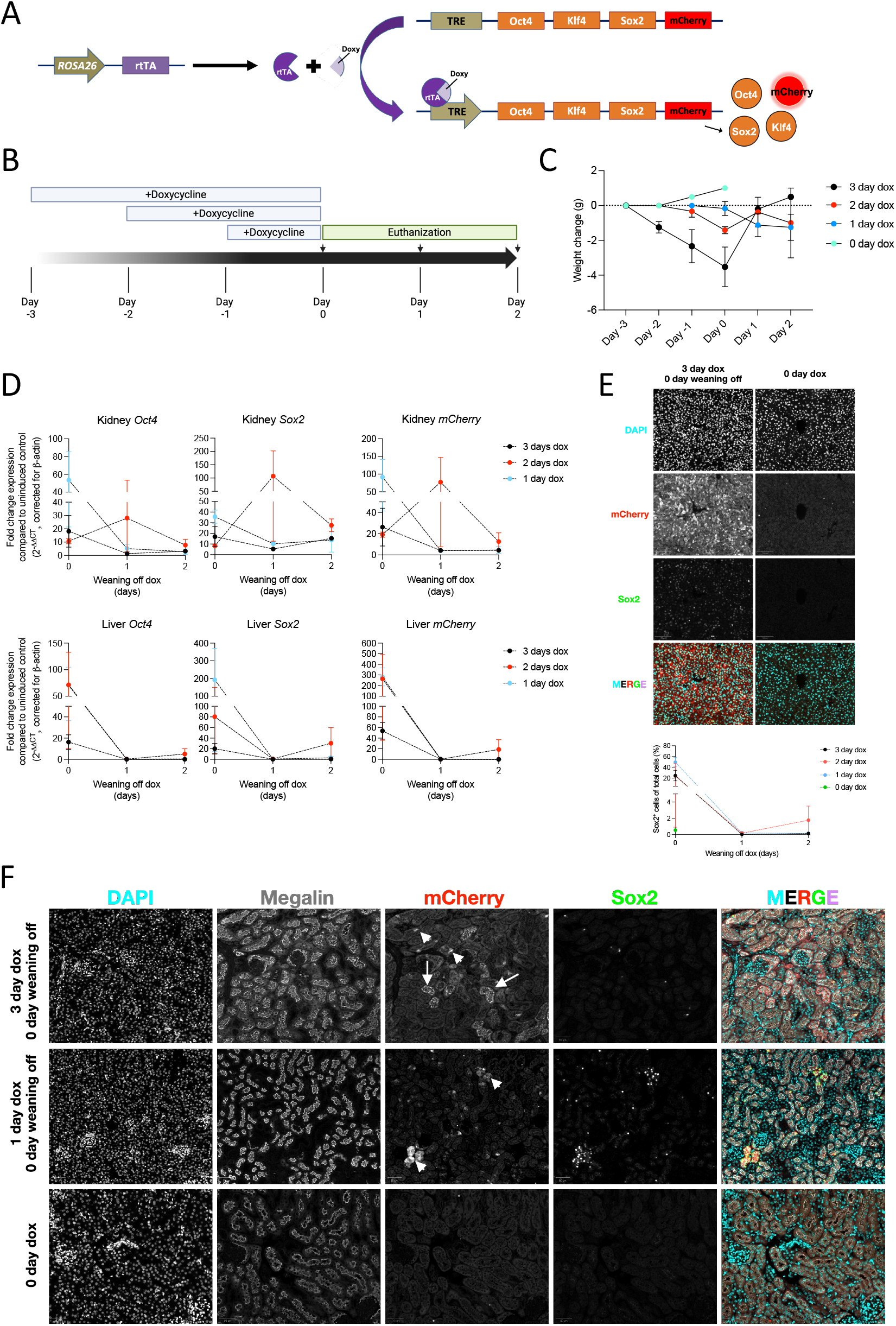
*ROSA26-*driven OKS is prominently expressed in liver, but not in kidney. A) Illustration of the gene structure used to induce the *OKSmC* ubiquitously. The rtTA is ubiquitously expressed and is activated upon binding to dox. Subsequent binding to the TRE leads to expression of the polycistronic *OKSmC* transgenes. B) Illustration of the timeline of the experiment performed with the tetO-*OKSmCherry*^tg/-^, ROSA26-*rtTA*^tg/-^ mice. Mice were exposed for different durations to dox and sacrificed at different time points. C) Weight change over time upon exposure to dox *ad libitum*. Each point is the mean and SEM of six mice, apart from uninduced control (N=1). D) Gene expression of *Oct4, Sox2* and *mCherry* in liver and kidney, showing mean and SEM of two biological duplicates, apart from groups where mice died as described. E) Immunofluorescence of the liver stained for mCherry and SOX2 and the quantification of SOX2+ cells per DAPI, depicted below. Average and SEM of two biological duplicates are depicted, apart from groups where mice died as described. Scale bars□=□100□µm F) Immunofluorescence of megalin, mCherry, and SOX2 in kidney of mice exposed to dox for 3, 1, or 0 day and sacrificed immediately afterwards. White arrows depict mCherry signal closely co-localizing with megalin, whereas arrow heads depict cytosolic mCherry signal. Scale bars□=□50□µm.

To determine the duration of dox exposure needed to detect *OKS* and assess their stability, a pilot experiment study was performed in which double heterozygous mice (tetO-*OKSmCherry*^tg/-^, ROSA26-*rtTA*^tg/-^) were exposed to 1 mg/mL dox *ad libitum* in their drinking water up to three days and sacrificed either immediately, one, or two days after cessation of dox exposure (**Fig 1B**). Noticeably, mice lost weight upon dox exposure, particularly those that were exposed to dox for three days, but regained weight after discontinuation of dox (**Fig 1C**). However, there was a 50% mortality in the subgroups that were weaned off dox for one and two days when exposed to dox for three days.

At dissection, macroscopically, no differences were observed in kidney or liver size or appearance between induced and control animals. No prominent histological differences were observed between different groups (**Supplementary Fig. 1A&B**). To verify the expression of the *OKS* transgenes, RNA was extracted from liver and kidney and RT-qPCR was performed. Expression for transgenes *Oct4, Sox2*, and *mCherry* expression was high in liver and kidney when tissues were collected directly after dox cessation, independent of its duration (**Fig. 1D**). Yet, after 24 hours or more after conclusion of dox treatment, expression of the *OKS* transgenes was lost. Immunofluorescence staining of liver revealed up to 49.5% of hepatocytes were SOX2+ when sacrificed immediately, whilst waiting for 24 hours following collection yielded similar SOX2+ hepatocytes as in controls (∼0.5%, **Fig 1E**). Though detected renal SOX2 expression was highest upon immediate harvesting of the organs upon cessation of dox (**Fig 1D**), the maximum of SOX2+ cells detected were ∼0.72% in the overall kidney (**Fig 1F**). Co-staining with megalin revealed that the majority of the SOX2+ population was within the megalin+ tubules, 1.12% vs 0.22% in megalin-cells.

Interestingly, when staining for mCherry, we observed two different signal patterns in proximal tubules, one being apically located in epithelial cells, and a second, more cytosolically located signal (**Fig. 1F**). Interestingly, only in the cells that exhibited cytosolic expression of mCherry, nuclear SOX2 expression could be detected. Moreover, kidneys from mice that received dox for 1 day did not show megalin+ cells with the apical mCherry pattern.

The observed low kidney expression, compared to the several fold higher liver expression, combined with the deteriorating health of mice after induction—thereby limiting the options for increasing dose or duration—prompted the conclusion that this ubiquitously expressing mouse model was not suitable for the planned study.

### Pax8Cre^+^ mice express *OKS* in proximal tubules

To determine if the transgenes could be expressed at a higher rate without inducing systemic toxicity, we next studied a kidney-specific expression mouse model. We used triple transgenic mice (tetO-*OKSmCherry*^tg/-^; *rtTA*^fl/-^; *Pax8Cre*^tg/wt^, hereafter Pax8Cre^+^ mice) that expressed Pax8-dependent rtTA (**Fig. 2A**). We administered dox in drinking water over up to 7 days and did not observe any weight changes or overt clinical impairment (**Fig. 2B-C**). Upon dissection, we observed no differences in respect of kidney size or appearance (data not shown). Periodic Acid-Schiff (PAS) and Masson’s Trichrome (MT) stainings of induced kidneys were similarly unremarkable (**Fig. 2D**). In expression analysis, we observed expression of the *OKS* and *mCherry* transgenes, with increasing fold-change until day 7 (**Fig. 2E**). This finding was confirmed using IF stainings for SOX2 and mCherry, with the highest expression detectable on day 7 (**Fig. 2F**). Of note, the increase in fluorescence signal for mCherry was more pronounced than for SOX2 although there was high colocalization between mCherry and SOX2 (**Fig. 2F**).

**Figure 2:**
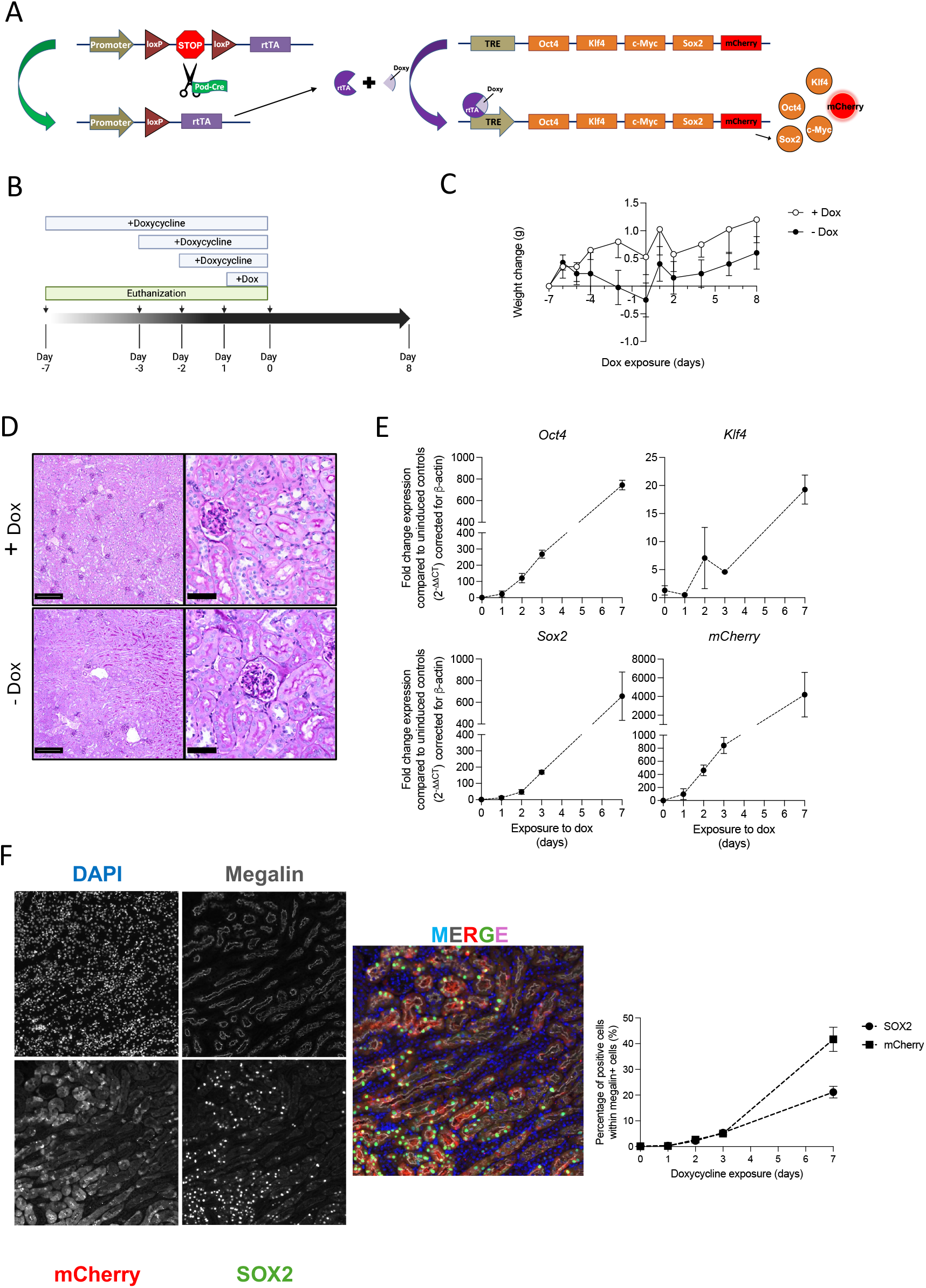
*Pax8-*driven OKS allows exclusive expression in kidney. A) Illustration of the gene structure used to induce the *OKSmC* in kidneys only. The rtTA is expressed in cells expressing Pax8-driven Cre recombinase and is activated upon binding to dox. Subsequent binding to the TRE leads to expression of the polycistronic *OKSmC* transgenes. B) Illustration of the time line of the experiment performed with the Pax8Cre^+^ and Pax8Cre^-^ mice. Mice were exposed for different durations to + or -dox and sacrificed at different time points. C) Weight change over time upon exposure to dox *ad libitum*. Each point is the mean and SEM of six mice, apart from uninduced control (N=2). No group effect or interaction effect was significant upon linear mixed-effect analysis. D) Representative images of PAS stainings of the kidneys of mice exposed for 7 days to + or -dox E) Gene expression of *Oct4, Klf4, Sox2* and *mCherry* in kidney of mice exposed to dox compared to -dox, showing mean and SEM (N=2). F) Immunofluorescence staining of SOX2 and mCherry in kidney of mice (left panel) exposed to dox for 7 days and counterstained for megalin with its quantification (right panel).

### Yamanaka factors induce dysplastic tumor formation in the kidney, dependent on induction length

Because of the known risk of tumor formation after induction with YF, we tested a 7-day induction protocol and waited for an additional 15 days before sacrificing the animals (**Fig. 3A**). Interestingly, we found multiple dysplastic tumors in both kidneys of all induced animals (**Fig. 3B**). Hence, we conducted additional experiments with lower duration of *OKS* induction and varying dox doses and waited for 4 weeks before sacrificing, to determine a safe induction protocol that would avoid tumor formation (**Fig. 3C**). One mouse in the 5 day-1 mg/mL group (N=3) developed tumors and cysts-like structures after a month (**Fig. 3D**), whereas both mice in the 5 day-2 mg/mL group developed tumors. However, we did not observe any dysplastic lesion in the 3-day long induced animals, for neither dose of dox (**Fig. 3D**). However, to minimize tumor risk, we opted for the lower dose of 1 mg/ml in our injury experiment.

**Figure 3:**
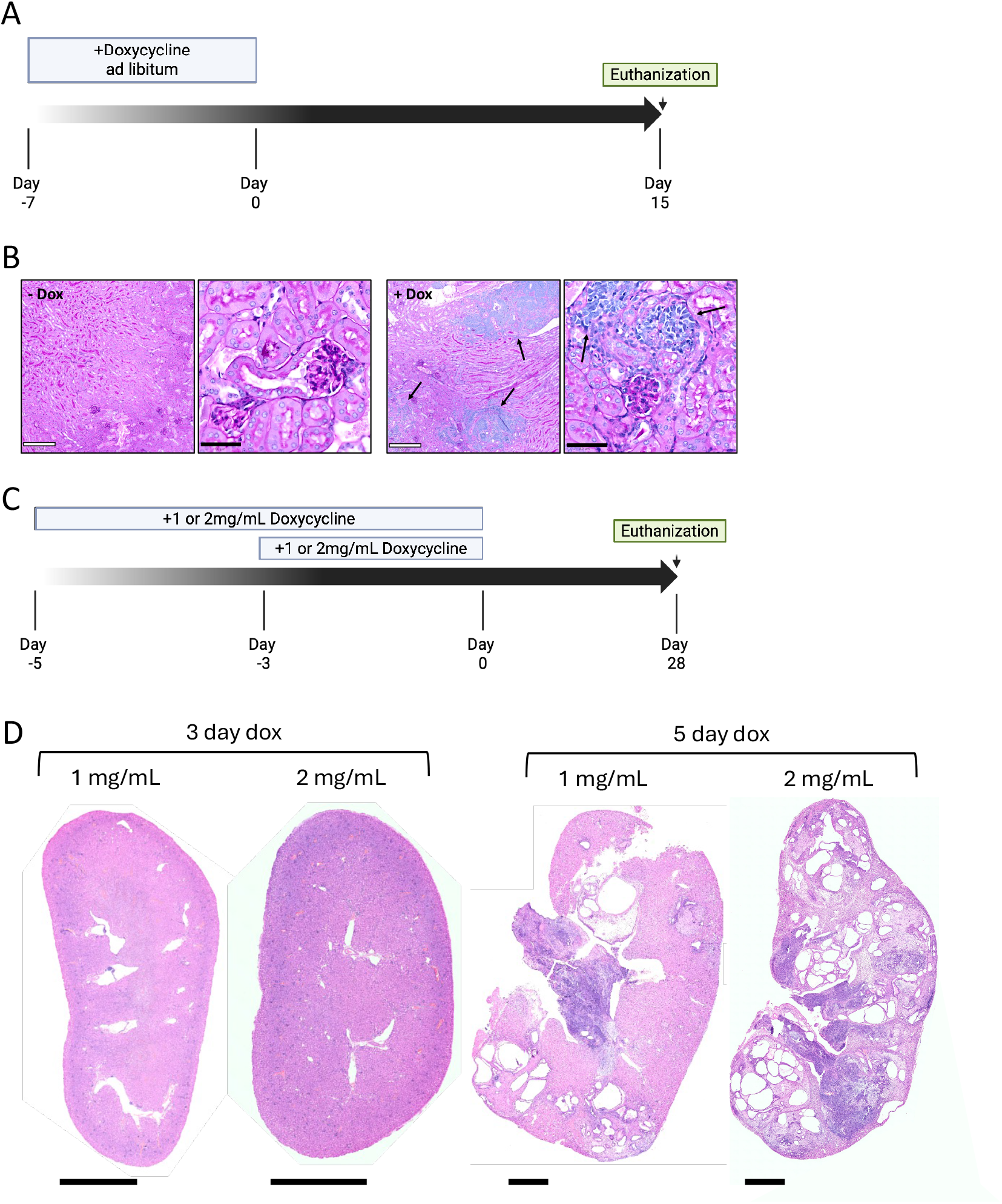
Relative long expression of *OKS* is associated with tumor formation. A) Illustration of the timeline of the experiment performed with the Pax8Cre^+^ and Pax8Cre^-^ mice. Mice were exposed for 7 days to + or -dox and sacrificed after 15 days (N=4 per group). B) Representative images of kidneys form mice exposed to + or -dox. Arrows point toward dysplastic tissues within the kidney (N=4 per group). C) Illustration of the timeline of the experiment performed with the Pax8Cre^+^ and Pax8Cre^-^ mice. Mice were exposed for to 1 or 2 mg/mL dox for 3 or 5 days. Mice were sacrificed after one month (N=2-3 per group). D) Images of whole kidneys post MT staining of mice exposed to different doses and duration of dox (N=2-3 per group).

### Yamanaka induction in kidneys did not ameliorate induced kidney injury

To investigate whether *OKS* induction could ameliorate induced kidney injury, we injected a dose of 7.5 mg/kg AA. Only male mice were used because significant sex differences for toxicity after AA administration had been established before.^29^ Since most *in vivo* studies on using *OKS* induction to ameliorate injury responses reported phenotypic amelioration when inducing *OKS* expression prior to injury,^14,21^ we decided to use the following protocol: mice were exposed to dox (1 mg/ml) for 2 days, following a single i.p. injection with AA or vehicle and continued dox induction for one more day, amounting to three days of *OKS*-induction in total (**Fig. 4A and B**). Both Pax8Cre^+^ and Pax8Cre^-^ mice receiving AA did not gain weight in contrast to vehicle-injected mice (**Fig. 4C**). When performing a grouped analysis based on whether animals received AA, this effect was significant using a linear mixed-effect model (AA:Genotype, p < 0.001, F: 34), however, there was no statistical difference between the Pax8Cre^+^ and Pax8Cre^-^ mice. Following dissection, blood analysis showed significantly increased levels of BUN upon AA administration, independent of genotype (**Fig. 4D**). To determine the long-term effects of transient *OKS* induction and the potential effects post AA-administration, we performed gene expression studies (**Fig. 4E**). Firstly, the *OKS* transgenes could not be detected in the Pax8Cre^+^ mice after the end point of the experiment (data not shown). We found *Havcr1*, encoding the proximal tubular kidney injury marker KIM-1, significantly increased in both Pax8Cre^+^ and Pax8Cre^-^ groups receiving AA compared to non-injured groups (p<0.001). Similarly, the AA-treated groups also showed a significant main effect on the distal injury marker *Lcn2* (NGAL; p<0.05), though no statistical difference was observed between individual groups. Notably, when comparing only vehicle-treated Pax8Cre^+^ and Pax8Cre^-^ mice using an unpaired Student’s T-test, the Pax8Cre^+^ mice showed significantly increased expression of these injury markers. In addition, the senescence markers *p16* and *p21* (p<0.05 and p<0.0001) were also increased upon AA-administration. Regarding fibrosis markers, a group effect was apparent between injured and non-injured mice for *Fn1* (p<0.01), but not for *Col4a1*, remarkably. *Tnf*α, an inflammation marker, was elevated in the injured groups (p<0.01), compared to non-injured groups. Importantly, in none of the tests a statistically significant difference was observed for Pax8Cre^+^ vs Pax8Cre^-^ groups (**Fig. 4E**).

**Figure 4:**
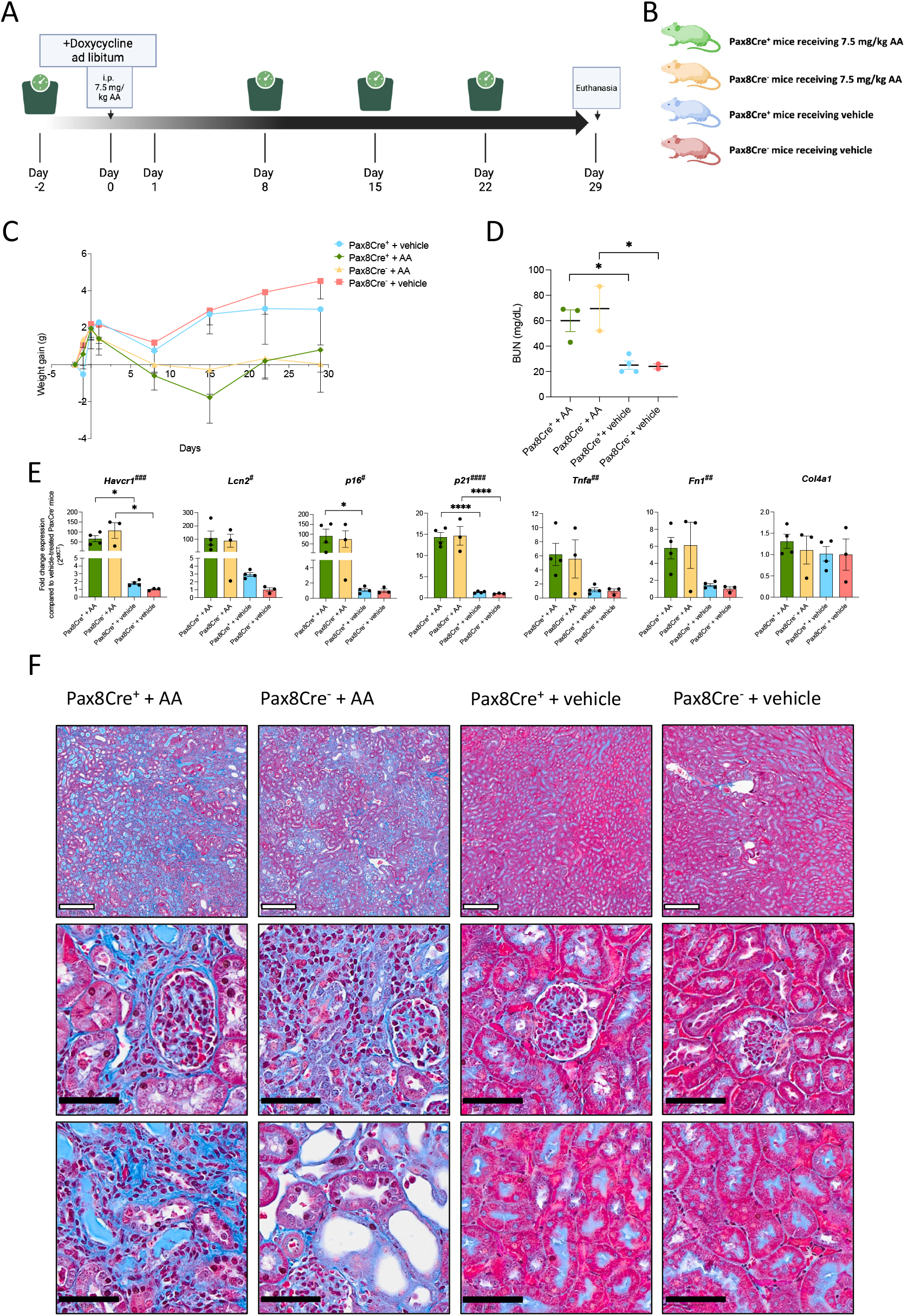
Single AA administration induces kidney injury, yet transient OKS expression does not yield amelioration of the phenotype. A and B) Illustration of the timeline of the experiment and the groups used for the injury experiment, respectively. Dox was given ad libitum on day -2 until day 1. 7.5 mg/kg AA or vehicle was i.p. administered on day 0. Mice were weighed at different time points and sacrificed after one month post dox. treatment. C) Weight changes across the experimental time. Each data point is the mean and SEM of N=3-4 mice. When comparing AA vs. no AA, linear mixed-effect analysis showed a significant interaction effect (AA:genotype, p < 0.001, F: 34), but no difference between YF induction / no YF induction. D) Blood urea nitrogen (BUN) at sacrificing time point. Depicted is the mean and SEM of N=2-4 mice per group. E) Gene expression of senescence, fibrosis, inflammation and injury markers. Depicted is the mean and SEM of N=3-4 mice per group relative to vehicle-treated Pax8Cre^-^ mice. Two-way ANOVA with Tukey post-hoc was performed. *P<0.05, between two individual groups, #P<0.05, ##P<0.01, ###P<0.001, ####P<0.0001 for overall AA-treatment vs vehicle F) Kidney, Masson’s Trichrome staining. White scale bar: 250 µm, black scale bar: 50 µm.

Histologically, the MT stainings of the kidneys revealed increased fibrosis of the kidney, particularly in the cortex (**Figure 4F**). Correspondingly, no differences using YF-induction were evident.

## DISCUSSION

We investigated two mouse models for *in vivo* induction of 3 Yamanaka factors, and performed an acute kidney injury experiment, testing the effect of YF on injury severity and outcome.

First, we used a *ROSA26*-driven ubiquitous *OKS*-expression mouse model, where we showed a high amount of transgene expression upon dox induction in hepatocytes (nearly 50% positivity) in immunofluorescence, but only a minimal induction of < 1% of kidney cells. Interestingly, there was an approximate 5-fold expression in megalin+ cells compared to megalin-cells. When analyzing mCherry specifically, we observed that there were more mCherry+ proximal tubules in contrast to SOX2+ proximal tubule cells. However, the majority of the mCherry signal was apically located, likely to originate from tubular uptake. As fluorescent reporter proteins are able to pass the glomerular filter,^30^ the observed apically located mCherry could stem from proteins that were synthesized and excreted by other organs such as the liver. This interpretation is further substantiated by the observation that mice exposed to dox for one day did not exhibit apical mCherry signal in the megalin+ cells, assuming that circulating amount of mCherry protein was likely not sufficient to be filtered and reabsorbed in the proximal tubule in a detectable manner.

Importantly, we observed a rapid decline in health and increased mortality upon dox induction of more than 2 days. This might have been due to previously reported liver or intestinal dysfunction upon induction.^31^ Taking this into account, we concluded that performing our injury studies in this mouse model was not promising and overall unethical.

Therefore, we pivoted to using a kidney-specific *OKS*-expressing mouse model. Pax8Cre^+^ mice exposed to dox showed a substantially higher gene expression of the *OKSmC* transgenes compared to our previous mouse model. Longer exposure to dox was also associated with higher expression of the transgenes. Moreover, no acute adverse effects were observed, allowing us to characterize this mouse model in greater detail.

Surprisingly, SOX2 and mCherry expression was observed almost exclusively in the proximal tubules in Pax8Cre^+^ mice, similar to the *ROSA26* mouse model, contrary to our hypothesis that in both transgenic mouse models all kidney tubular cells would express the *OKSmC* transgenes. Both *ROSA26* and *Pax8* promoters are active throughout different segments of the kidney with several studies describing transgenic expression in proximal and distal tubules.^32–34^ Yet, the vast majority of SOX2 expression was detected in proximal tubules. Of note, it is possible that the immunofluorescences technique used may not be sensitive enough to detect more moderate expression in other cell types.

We chose the aristolochic acid injury mouse model, since it is well established as a model to study AKI to CKD transition, with progressive tubulointerstitial nephritis. In fact, AA has been shown to primarily injure proximal tubules,^35^ matching the YF-expressing location. However, for the here studied timeline and YF induction regime, we did not observe clinical differences when analyzing histology and blood parameters, or injury markers on transcriptional level -in contrast to various previous studies which showed an amelioration of injury after *in vivo* induction with YF in multiple organ systems.

So far, no injury study in conjunction with YF application targeting the kidneys has been published, but two studies investigating age-related changes found significant amelioration of aging phenotypes on molecular and histological level in the kidney,^11,19^ with tubular atrophy in their transgenic progeria mice, which was not evident in the treated mice. In addition, previous studies using kidney progenitor cells injected into living mice, involving cisplatin-induced and AA-induced AKI and CKD, could show integration of cells in proximal renal tubules, with significant protective effects on histologic and clinical phenotypes. ^36,37^ Two other studies used iPSCs for direct injection into rats, demonstrating significant effects on both AKI and CKD, albeit with the risk of tumor formation in the kidney in the CKD study.^38,39^ Taken together, these studies showed promising potential for inducing a kidney progenitor state in a kidney injury model to prevent CKD progression and ameliorate kidney function.

However, there are several possible explanations which could have hampered improved regeneration in our study.

First, we were only able to show expression in approximately a quarter of proximal tubular cells, which could have limited the potential therapeutic effect. Second, it is well established that after acute injury not only tubular cells contribute to transition to kidney fibrosis and CKD, but also endothelial cells in the interstitium, migrating inflammatory cells, as well as myofibroblasts.^40^ Possibly, induction of YF limited to proximal tubular cells cannot compensate for detrimental effects caused by the other cell types. To elucidate this aspect, an additional mouse model with whole kidney YF expression would be required. Third, the observed injury was relatively strong, with substantial fibrosis after 4 weeks. Potentially, a milder injury would have been more responsive to amelioration by YF. Fourth, we were only able to expose mice to dox for a limited amount of time due to dysplastic transformation (as discussed in more detail in the next paragraph), pointing towards an especially high sensitivity of proximal tubular cells regarding loss of identity and full reprogramming.

### Partial reprogramming and cancer formation

When we conducted induction experiments in the kidney-specific YF-expressing mouse model, we surprisingly observed vast tumor formation after relatively short induction periods despite omission of the highly cancerogenic *m-Myc*. Interestingly, when Onishi et al. investigated dysplastic tumors that were forming after YF induction in various organs using a mouse model with ubiquitous expression of all four YF (OSKM), they concluded that these represent “failed reprogrammed cells”.^28^ They encountered dysplastic cells that would undergo redifferentiation to “normal” cells after dox withdrawal, however, if induction length reached a certain point (typically 3-4 weeks), the dysplastic cells would further proliferate and transform into cancer-like dysplastic tumors.^28^ Importantly, also in mice expressing only *OKS*, they did encounter these dysplastic tumors, albeit only after a prolonged induction time span of 21 days, so, even compared to this other study, tumor formation in our study was fast. Importantly, studies using a cycling YF induction (2 days or 3 days induction, followed by 5 or 4 days of no induction, respectively), showed anti-aging effects in a large variety of tissues, including kidney, but no formation of teratoma, even after more than 10 months of treatment in this cycling pattern.^11,17,19^ Another study on aged mice and a progeria mouse model could ameliorate age-related changes in multiple organs, again including kidney, using a continuous treatment of dox for 2.5 weeks.^18^ The authors did not observe any negative effects, albeit using a transgenic mouse model with all 4 YF, but using lower doses of dox in the drinking water.

We cannot exclude that using a cyclic induction pattern in our experimental setup would have yielded an improved outcome. Because of the singular injury event however, we did not consider a cycling induction especially intuitive. Additionally, also a lower dox dose might have achieved differential effects, in combination with possibly less dysplasia formation. Yet, this points towards an extremely narrow therapeutic window between achieving beneficial effects on one hand side and cancer formation on the other, especially in a tissue type that seems to be especially susceptible to dysplastic transformation after YF induction. Especially, since our transcriptional analysis of kidney injury markers indicates that YF induction alone already caused a limited amount of kidney injury.

## Conclusion

*In vivo* application of the Yamanaka factors to induce partial reprogramming has produced remarkable improvements in regeneration, cellular plasticity, and age-related phenotypes, particularly in the nervous system. However, tissues vary substantially in both their reprogramming capacity and susceptibility to oncogenic transformation, as demonstrated in kidney-specific models. Determining a dose that promotes repair without crossing the threshold into deleterious effects remains a major challenge, raising significant concerns about the near-term feasibility of translating this approach to humans.

## AUTHOR CONTRIBUTIONS

Conceptualization: K.L., G.F., F.B., F.H.; Data curation: K.L., G.F.; Formal analysis: K.L., G.F.; Funding Acquisition: F.H.; Investigation: K.L., G.F., S.H., K.M.R., K.Y., C.M.K., E.Z., D.M., K.S., Kr.L., G.K.; Methodology: K.L., G.F., Project administration: F.H.; Supervision: F.B.; Visualization: G.F., Writing – original draft: K.L., G.F.; Writing – review and editing: K.L., G.F., F.B., F.H.

## FUNDING

F.H. is the William E. Harmon Professor of Pediatrics at Harvard Medical School. This research was supported by grants from the National Institutes of Health to F.H. (RC-2-DK1222397), to K.Y (3T32DK007726-38S1), the German Research Foundation to K.L. (Project No.: 461126211), to C.M.K. (Project No.: 499462148 and DFG-Rückkehrstipendium (KO 6579/ /3-1), to F.B (Project No.: 404527522), and to K.R (Project No.: 519309154), the JSPS Overseas Research Fellowship to K.S. (No. 202260295), the German Academic Exchange Service via the Biomedical Education Program to S.H., the iPRIME Clinician Scientist Forschungskolleg (2021_EKFK.15, UKE, Hamburg, Germany) from the Else Kröner-Fresenius-Stiftung to FB, the Carl W. Gottschalk Research Scholar Grant from the American Society of Nephrology to F.B, and the Netherlands Organization for Scientific Research (Rubicon: 452022311) to G.F.

**Supplementary Figure 1: *ROSA26*-driven OKS expression does not lead to histological changes in kidney or liver**.A) Representative H&E staining of kidney of mice, immediately collected after 3-day exposure to dox or control. White scale bar: 200 um, black scale bar: 50 µm. B) Representative H&E staining of liver of mice, immediately collected after 3-day exposure to dox or control. White scale bar: 200 um, black scale bar: 50 µm.

