## Supplementary figures and images for "A kidney specific mouse model to study the effects of *in vivo* induction of Yamanaka factors"

### Supplementary Figure 1

#### Supplementary figure 1

A

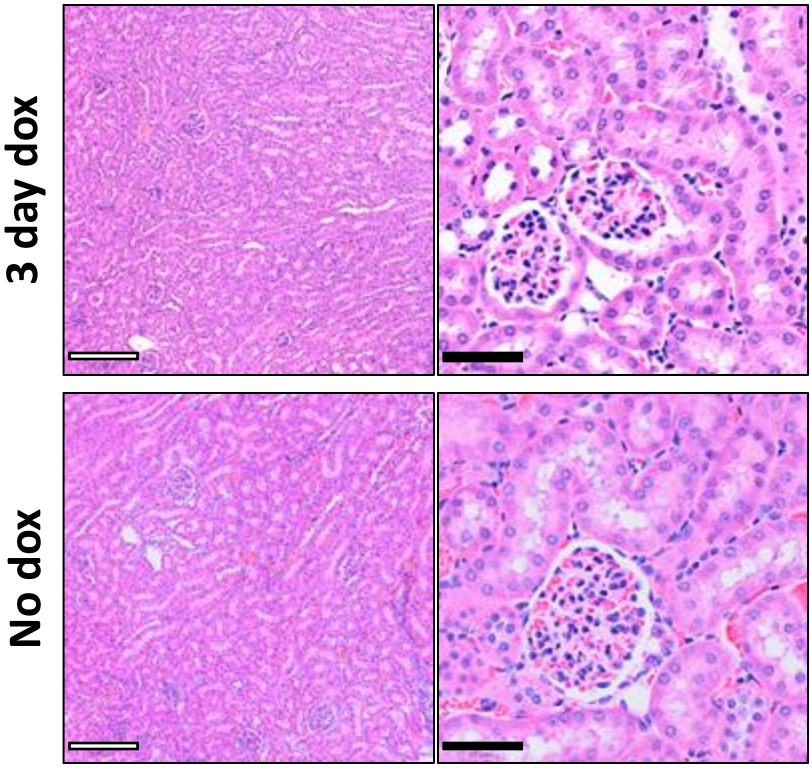


B

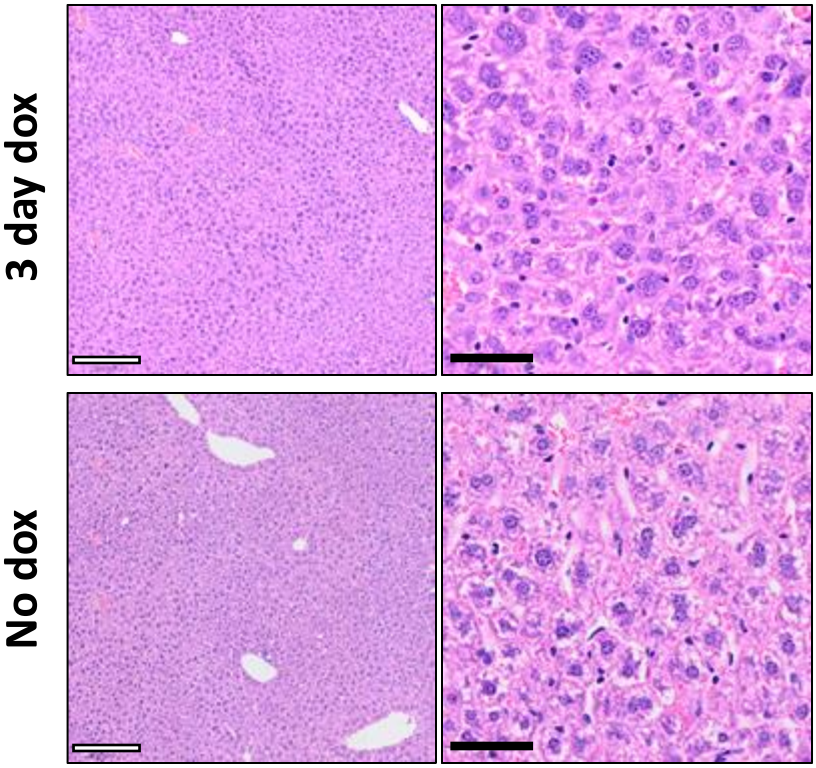
